# Climatic niches in phylogenetic comparative studies: a review of challenges and approaches

**DOI:** 10.1101/018796

**Authors:** Lara Budic, Carsten F. Dormann

**Affiliations:** Faculty of Environment and Natural Resources, University of Freiburg, Tennenbacher Str. 4, 79106 Freiburg

**Keywords:** Climatic niche, phylogenetic comparative studies, ancestral niche reconstruction, evolutionary models

## Abstract

Studying the evolution of climatic niches through time in a phylogenetic comparative framework combines species distribution modeling with phylogenies. Phylogenetic comparative studies aid the understanding of the evolution of species’ environmental preferences by revealing the underlying evolutionary processes and causes, detecting the differences among groups of species or relative to evolutionary pattern of other phenotypic traits, but also act as a yardstick to gauge the adaptational potential under climate change. Because several alternatives exist on how to compute and represent the climatic niche, we here review and discuss the current state of the art and propose a best practice to use in comparative studies. Moreover we outline the common evolutionary models and available model-fitting methods and describe the procedure for ancestral niche reconstruction with the intention to give a broad overview and highlight the most advanced approaches for optimal niche-related comparative studies.

## Introduction

Phylogenetic comparative studies use a wide range of methods to explore patterns and processes linked to phylogenetic trees and species traits (Pennell and Harmon, 2013). These studies uncover how a certain trait evolves among different taxa, how evolution of one trait influences another, whether a trait represents adaptation to the environment etc. In this review we focus exclusively on studies testing hypotheses about species’ climatic niches evolution through phylogeny.

The aim of such studies is typically not only to suggest the trajectories of niche evolution, but rather to test specific hypothesis about the timing of appearance, causation or evolutionary processes responsible for observed patterns. Such studies aim to discover, for instance, whether shifts in the climate niche occur at the same time as shifts in a particular trait (such as C3/C4 photosynthesis: Edwards and Smith, 2010), whether it was a key driver for developing specific life-histories (e.g. cactus life-form: Edwards and Donoghue, 2006), or whether temporal and/or spatial fluctuation of climate caused species to evolve and diversify (Evans et al., 2009). Furthermore, linking climate niche evolution to population demography through time could reveal whether major niche shifts occur in small or large populations (Jakob et al., 2010). The analyses are based on current trait values, the phylogenetic relationship between species and an evolutionary model. Climate niche is treated as if it was a phenotypic trait and the analysis of, say, temperature values follows the same logic as evolutionary analysis of body mass. The actual reconstruction of ancestral trait values is unnecessary for testing correlations between characters. Being an integral part of the procedure, ancestral states along the phylogeny are implicitly inferred but not actually presented. However, other hypotheses might require reconstruction of ancestral values, such as tests of niche overlap between specific ancestral nodes, pinpointing the exact time of appearance of certain values, identifying reversals in evolutionary progression or visualization of trait changes through time.

Given that understanding the evolution of species’ niches through time attracts extensive interest and is highlighted as a priority research question in paleoecology (Seddon et al., 2014), we discuss and propose guidelines for optimal use of species distribution and climatic data in comparative studies.

The aim of this review is to (1) introduce the concepts relating niche space to its evolution through time, (2) discuss and propose the optimal niche representation to be used in phylogenetic comparative studies, (3) introduce most common evolutionary models (4) describe the methods for ancestral reconstruction and discuss future directions of the field.

### Climate niche in space and time

Hutchinson (1957, 1978) defined a species’ fundamental niche as all combinations of environmental conditions where a species can persist and maintain a viable population in the absence of predators or competitors (Kearney and Porter, 2004). Although the ecological niche is not strictly a heritable phenotypic trait for which these methods were developed, niche characteristics are defined and constrained by species physiology, which *is* heritable, and as such can be analyzed in a phylogenetic framework (Kozak and Wiens, 2010a).

Biologists are generally interested in the “fundamental niche”, which represents species physiological limits and is the actual evolvable trait, although what we actually observe in nature is the “realized niche”. Unfortunately we are currently unable to determine the fundamental niche without manipulative experimental studies, so we are bound to analyse the realized niche, a restricted section where a species lives, limited by biotic interaction or dispersal limitation (Jackson and Overpeck, 2000; Soberón and Nakamura, 2009).

It is currently impossible to tell how closely the realized niche approximates the fundamental niche, and for this reason it is difficult to guess whether the observed change really demonstrates niche evolution, or if this change is merely a shift of the realized niche within the species’ fundamental niche (e.g. due to changed biotic interactions: Graham et al., 2004; Dormann et al., 2010). High within-species plasticity (e.g. in mammals, Réale et al., 2003) may lead to changes in realised habitats as biotic and/or abiotic conditions change.

A further complication is introduced by the existence of no-analog climate conditions at different time slices, e.g. in the past (Williams et al., 2001) or future (Williams et al., 2007, see Fig.1), which indicates that only a portion of a species’ fundamental climatic niche (termed “potential niche”) actually exists in the world at a given time. Therefore, whole sections of fundamental niche might be unobserved because they are nonexistent in space at that time (Fig. 1). Moreover, in the presence of facilitative biotic interactions or mutualism, species’ niche could even extend beyond the fundamental niche (e.g. in lichens or corals; see Fig. 1, Bruno et al., 2003; Afkhami et al., 2014). Given the tremendous complexity of confounding factors it seems incredibly difficult to be certain how to interpret the observed change in the climate niche. Distinguishing evolutionary change from shifts within unknown niche limits certainly merits attention, and comparative studies could potentially disentangle one from another: if a trait tightly linked to physiology changes along with the niche (assuming this change is at the genetic level, i.e. it goes beyond phenotypic plasticity), this could indicate the species is adapting to new environmental conditions, and hence its niche is evolving. For example, Edwards and Smith (2010) found that the origin of C4 photosynthesis in grasses coincided with shifts to drier environments. Without considering the exact mechanism, or excluding other hypotheses, it should be reasonable to interpret this finding as species adaptation to novel climate conditions (i.e. evolutionary change of the fundamental niche). In this review we consider the “realized” niche, since this is the most common situation for which we have data although we acknowledge it is far from a consistent approximation of the fundamental niche, and encourage using direct physiological estimates of climatic tolerance whenever available.

**Figure 1:**
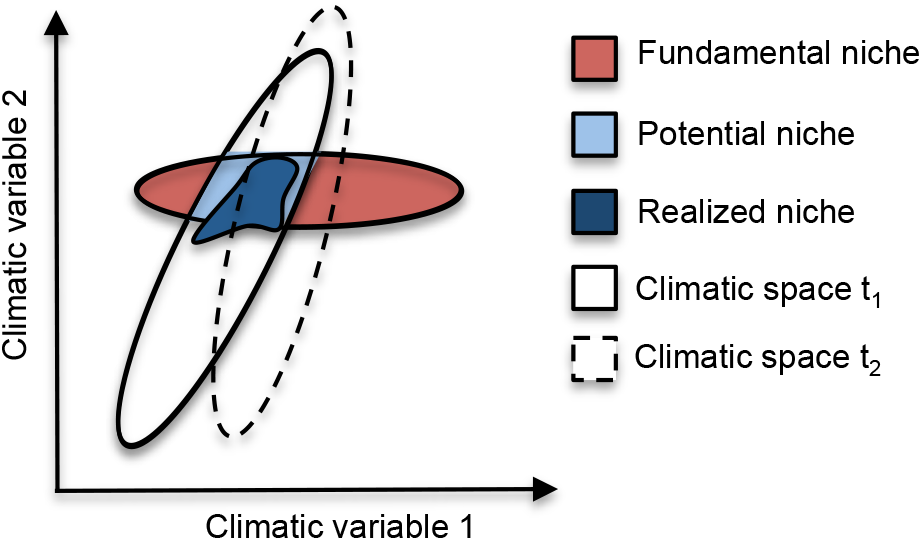
Climatic space represents all the combinations of two climatic variables existing at a certain time, which may differ at different times in history (t1, t2). The fundamental niche of a species includes all the possible conditions where a species could persist, some of which may lie outside conditions currently existing in the world. The intersection of the two represents the potential niche, which the species would fill in the absence of biotic interactions and dispersal limitations. The realized niche is the segment actually occupied by the species; it may occasionally extend towards climatic conditions outside its fundamental niche if facilitative biotic interactions are present. Adapted from Jackson and Overpeck (2000).

**Figure 2:**
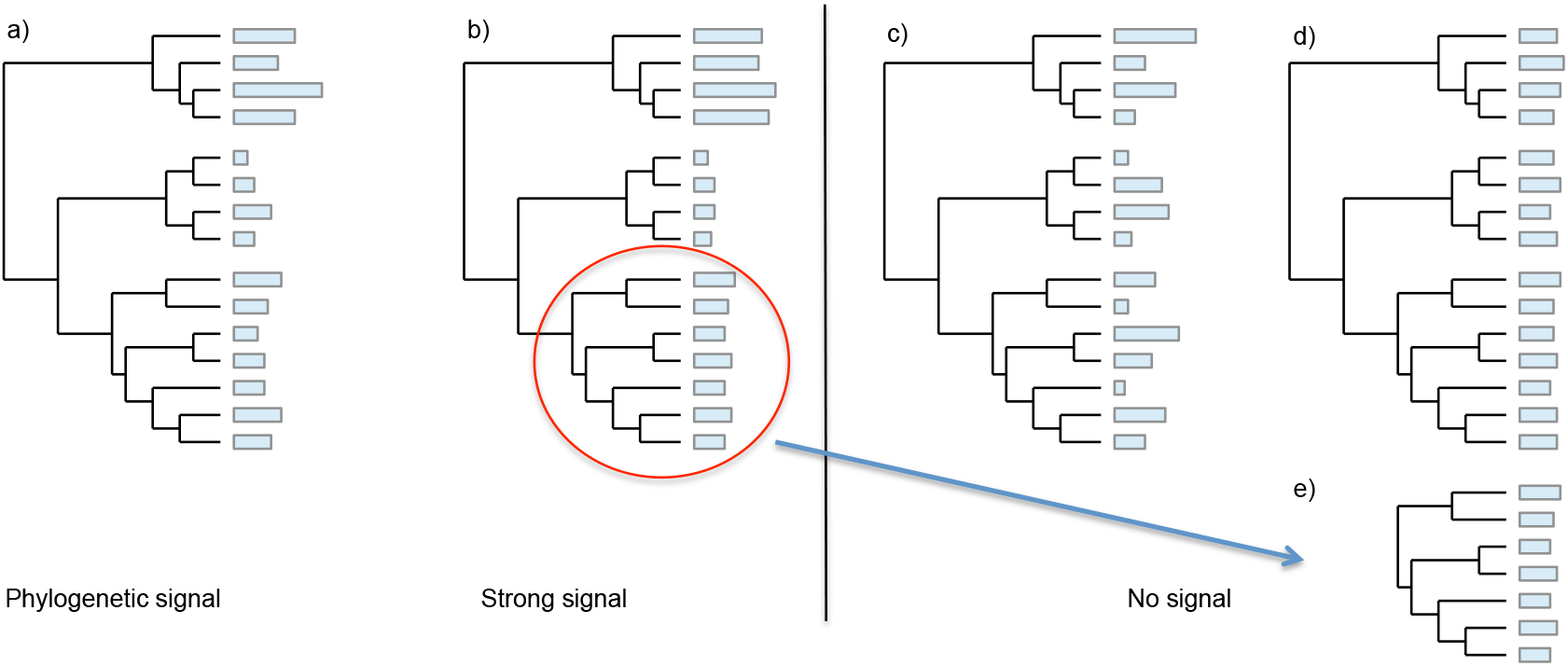
A hypothetical illustration of different degrees of phylogenetic signal. (a) Situation where trait values follow the expectations of a BM model of evolution (i.e. traits similarity is proportional to shared evolutionary history among species). (b) Strong phylogenetic signal is found when the trait values among closely related species are more similar than would be expected from BM. (c) A situation with no or low phylogenetic signal as a consequence of over-dispersion or convergence to habitat-specific trait values. Closely related species are not more similar than species drawn at random. (d) No phylogenetic signal as a consequence of trait stasis or convergence of all the species to the same trait value. (e) A signal (or its absence) may be restricted to a specific clade of a phylogenetic tree.

### Representing the climatic niche

To test hypotheses about the climatic niche in phylogenetic comparative methods, it is necessary to infer the present climate niche of extant species. There is no standard protocol and the way the niche is represented varies widely among studies. For example, some authors represent the niche with climate niche models (Graham et al., 2004; Yesson and Culham, 2006a,b; Eaton et al., 2008; Dormann et al., 2010), others use raw climate data (Evans et al., 2005; Ackerly et al., 2006), some combine the two approaches (Fig. 3, Evans et al., 2009; Smith and Donoghue, 2010), and others represent the niche with ordination techniques (principal component analysis, outlying mean index: Eaton et al., 2008; Boucher et al., 2012; Bystriakova et al., 2011). Here we define the optimal climate niche representation for the most common circumstance, where data derive from occurrence data and the hypothesis to be tested is specific to that region. In this case, niches based on **raw** data consist only of climate values extracted at species locations: generally the mean of those values is representing the niche (e.g. mean annual temperature, total annual precipitation). Climate niche models, a subset of species distribution models, are the most frequently used approach to represent the niche (Franklin and Miller, 2009; Peterson et al., 2011). Algorithms relate a species’ geographical locations to climate characteristics in order to describe its environmental niche (Guisan and Zimmermann, 2000; Kozak et al., 2008, a review of prediction ability of numerous modeling algorithms is provided by Elith et al. 2006). Modeling the niche to get insights of the functional relationship between a species and its environment is statistically preferable to the use of raw data (Peterson et al., 2011), because it accounts for the fact that species occurrences in an area might be determined by habitat availability and is not only a function of species preferences. For instance, a higher abundance in valleys compared to mountain tops could be due to a higher availability of valleys in that area, despite species’ higher preference for mountains (see Fig. 3 c,d and examples below).

**Figure 3:**
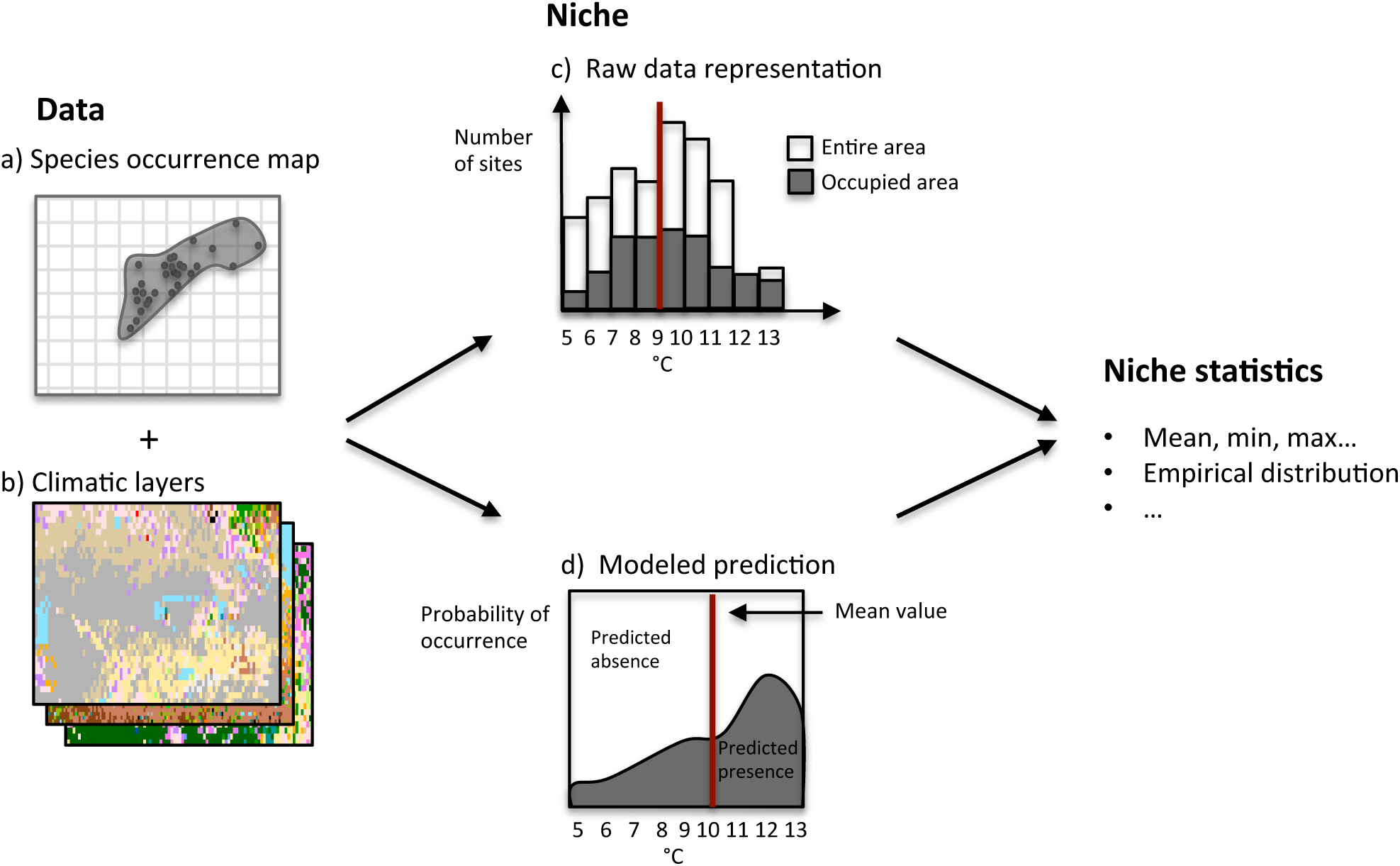
Climate niche representation. Niches are inferred by combining a) species distribution maps with b) climatic layers (temperature, precipitation, etc). c) The simplest niche representation that combines those data can be plotted as a histogram, with grey bars being the number of species occurrences at a certain temperature and white bars representing the entire area. Alternatively, niches can be statistically modeled, as shown in diagram d). The parameters finally obtained from those two approaches, in this example the mean values, may differ. Fig d) illustrates why the mean value is not necessarily the optimal statistic; instead reconstructing the whole distribution is preferable.

#### Niche statistics

The niche is a multidimensional entity and as such difficult to analyze phylogenetically as a whole. In general, it is decomposed into its marginal components (e.g. annual precipitation), each of which is examined separately along the phylogenetic tree. Most if not all variables describing the climate niche are continuous and the statistic most commonly chosen to represent them is the mean (e.g. mean annual precipitation). As some authors acknowledged, the mean may or may not be the most informative descriptor of the niche (see Fig. 3d, Graham et al., 2004). Different solutions have been proposed, particularly among studies where ancestral climate niche was reconstructed along the phylogeny. Graham et al. (2004) and Hardy and Linder (2005) proposed to consider the upper and lower niche limits separately, in order to infer the whole range of conditions of the ancestral niche, the so called “MaxMin” coding, which was used in a number of later studies (Yesson and Culham, 2006a,b; Lo Presti and Oberprieler, 2009; Lawing and Polly, 2011; Töpel et al., 2012). Instead of using maximum and minimum, which could be outliers, Vieites et al. (2009) proposed to consider 95% confidence values. In any case, mean, minimum or maximum temperature still do not fully describe the distribution of species climatic tolerances. To tackle this issue Evans et al. (2009) proposed the “predicted niche occupancy” (PNO) profiles: histograms obtained by combining response curves from niche models with climate layers of actual species distribution in geographic space. With this approach, each climate variable is represented by a histogram, which is especially appropriate for species whose niche variables are multimodal or, more generally, do not approximate a normal distribution. Working with histograms (or rather, empirical densities) requires sampling from the distribution of values and thus repeating the same analysis for each sampled value.

Analyzing the whole distribution of preferences is certainly more desirable than using single values. It would, however, be more appropriate to consider response curves obtained from niche models in parameter space, rather than combining them with geographic space. This avoids spurious results arising when large areas with low suitability are present in geographic space: the total sum of suitabilities (as used in Evans et al., 2009) could still be higher for those suboptimal conditions, only because of their high frequency in geographic space. This is illustrated in Fig. 3c, where a great portion of sites with the temperature of 9-10° C are not occupied by the species, therefore the probability of species occurrence at that temperature is relatively low (see Fig. 3d), but the sum of those probabilities in space will be high because there are many sites with that temperature. On the other hand, a temperature of 12-13°C is highly suitable (probability of 0.8), but there are so few sites with these conditions that the sum of suitabilities is low, despite the species’ high preference for those sites. The statistical model would pick it up and discriminate between use and availability (the maximum in Fig. 3d), while the histogram of suitability in geographic space (PNO) will be biased towards common environmental conditions. Therefore, model output allows an unbiased representation of preferences, irrespective of geographic availability (Hurlbert, 1978; Manly et al., 1993; Matthiopoulos et al., 2011). This is relevant also when considering large-scale climate change and the existence of no-analog climates in different time periods - climate conditions that cover large areas today might have been very restricted at a different time, and vice versa. In principle, the same resampling scheme can be used for multivariate distributions (as in Boucher et al., 2012), obviating the need to study each climate variable separately. However, given the no-analog conditions, the problem of assuming the same correlation structure between variables for different time periods arises again, an issue still waiting to be solved.

#### Ecological variability

Species are often polymorphic; populations of the same species may live, for example, on different types of soil, or along a wide gradient of temperature (Pearman et al., 2010). The approach described above, where species’ climatic preferences are expressed with empirical densities, automatically takes into account the ecological variability within species by resampling from climatic values based on species’ preferences. Theoretically the same approach could be employed for categorical variables, with preferences determining the probability of drawing from each character state (e.g. in case of higher species’ preference for soil-type A compared to B). This way ecological variability can be taken into account in any comparative analysis. Another method to accommodate polymorphism in discrete characters is through the quantitative genetic threshold model (Felsenstein, 2005), which models a discrete character as a continuous trait and is described in more detail in Box 1. See also Hardy and Linder (2005) and Hardy (2006) for additional methods and discussion on this topic.

### Evolutionary analyses

How did the niche evolve among different species? Under which processes? What were the drivers? Are niches conserved or labile? What was the ancestral niche like? These are some of the most intriguing questions in comparative studies about climatic niches. We next describe the available methods to tackle some of them: we discuss the utility of phylogenetic conservatism tests, describe the most common evolutionary models, explain the procedure to infer the ancestral climatic niche and end with a summary and recommendations of best approaches.

As the name suggests, phylogenetic tree is the backbone of phylogenetic comparative studies. A detailed description on building a phylogeny is beyond the scope of this review, and we refer the interested readers to Holder et al. (2003); Bininda-Emonds (2004) and Roquet et al. (2012). At the coarsest level, a distinction can be drawn between phylograms (trees with branch lengths proportional to molecular distance), and chronograms (or ultrametric trees, with branch lengths proportional to time). Unless the aim of the study specifically requires the use of a phylogram, the general consensus is to use ultrametric trees because niche evolution is assumed to be proportional to time. Although sophisticated methods are continually reducing uncertainty, phylogenies still remain only hypotheses of how species evolved (Webb et al., 2002). Alternative trees often have almost the same support, which is problematic because for instance, niche reconstruction on different trees may produce different results. Therefore the best way to incorporate phylogenetic uncertainty is to carry out the analyses on a sample of plausible phylogenetic trees instead of using the single best phylogeny.

#### Phylogenetic niche conservatism

Phylogenetic niche conservatism Phylogenetic niche conservatism (PNC) is the tendency of species to retain their ancestral niches through time (Boucher et al., 2014). The most common way to assess the PNC is by measuring the phylogenetic signal: a measure indicating whether a trait evolves according to the null expectation of neutral drift model. There is still disagreement and it remains a debated topic at which similarity level phylogenetic signal can be interpreted as phylogenetic niche conservatism (Losos, 2008a; Wiens et al., 2010).

Here we want to highlight how testing for PNC by measuring phylogenetic signal can be potentially misleading and special caution is needed when interpreting the results. For instance, no phylogenetic signal is a pattern where species niches appear to be independent from phylogenetic relationship among them (Losos, 2011). This is usually interpreted as no niche conservatism, as it can arise when the niche evolved more than expected from random evolution. Niche diverged to such an extent that the similarity among closely related species is lost (Fig. 1c). Another cause leading to the same pattern is convergence, when species belonging to separate clades adapt to the same types of environment, and therefore the pattern of niche values distribution among clades is similar (Fig. 1c, Kraft et al., 2007). Again, this is seen as no niche conservatism. But a highly problematic and less obvious cause of observing no phylogenetic signal is *perfect* conservatism: if the evolution is extremely conserved, all species will have the same or very similar niches, and no phylogenetic signal can be detected (Fig. 1d, Revell et al., 2008; Kozak and Wiens, 2010b). This occurs under strongly stabilizing selection, where all species evolve towards the same optimum value (Revell et al., 2008; Kozak and Wiens, 2010b), or when strong biological constraints bound the niche values to a narrow interval (Revell et al., 2008; Losos, 2011). Hence no signal could indicate either divergent evolution, strong stabilizing selection with one optimum (i.e. stasis), or bounded evolution. Therefore, the same pattern can be caused by completely different processes, which cannot be distinguished among each other by measuring the level of phylogenetic signal alone. To identify whether the niche evolved under, e.g., directional selection or genetic drift, the recommended approach is to fit different evolutionary models to data, rather than measuring the phylogenetic signal (Revell et al., 2008; Cooper et al., 2010).

Furthermore, the detection of a phylogenetic signal depends on the size of the phylogenetic tree and the section analyzed (Fig. 1e). It is extremely important to interpret the patterns only according to the climatic/temporal boundaries within which they were identified; niche lability in a strictly tropical species does not preclude PNC at larger scales (Losos, 2008b; Wiens, 2008).

To summarize: (a) phylogenetic signal and niche conservatism are patterns which do not necessarily reveal the underlying processes (Losos, 2011; Crisp and Cook, 2012); (b) completely different processes can lead to the same pattern (see Revell et al., 2008); (c) the detection of patterns is context-dependent (Fig. 1e).

Therefore, a better approach to assess niche conservatism among different clades is to test for mechanisms and evolutionary processes.

#### Models of evolution

Evolutionary models describe and approximate the natural processes responsible for trait evolution (Fig. 4). Fitting various models to the data permits to test hypotheses about the processes driving evolution of particular trait (e.g. the climatic niche). Before actually fitting models to the data, it is advisable to first identify plausible evolutionary processes based on prior biological knowledge, and afterwards proceed to fit only the corresponding models (Fig. 5). By fitting all the models without distinction one runs the risk of selecting a model with good statistical fit, but biologically improbable premises (as demonstrated in Wiens et al., 2007). The model with best fit is chosen in most cases through likelihood ratio tests (LRT, Johnson and Omland, 2004). Model selection criteria such as Akaike information criterion (AIC, Akaike, 1973) or Schwarz criterion (BIC, Schwarz, 1978) provide several advantages; they can compare multiple models simultaneously, rank them, give relative supports and are not influenced by the hierarchical order in which the models are compared (Burnham and Anderson, 2002; Johnson and Omland, 2004; Posada and Buckley, 2004).

**Figure 4:**
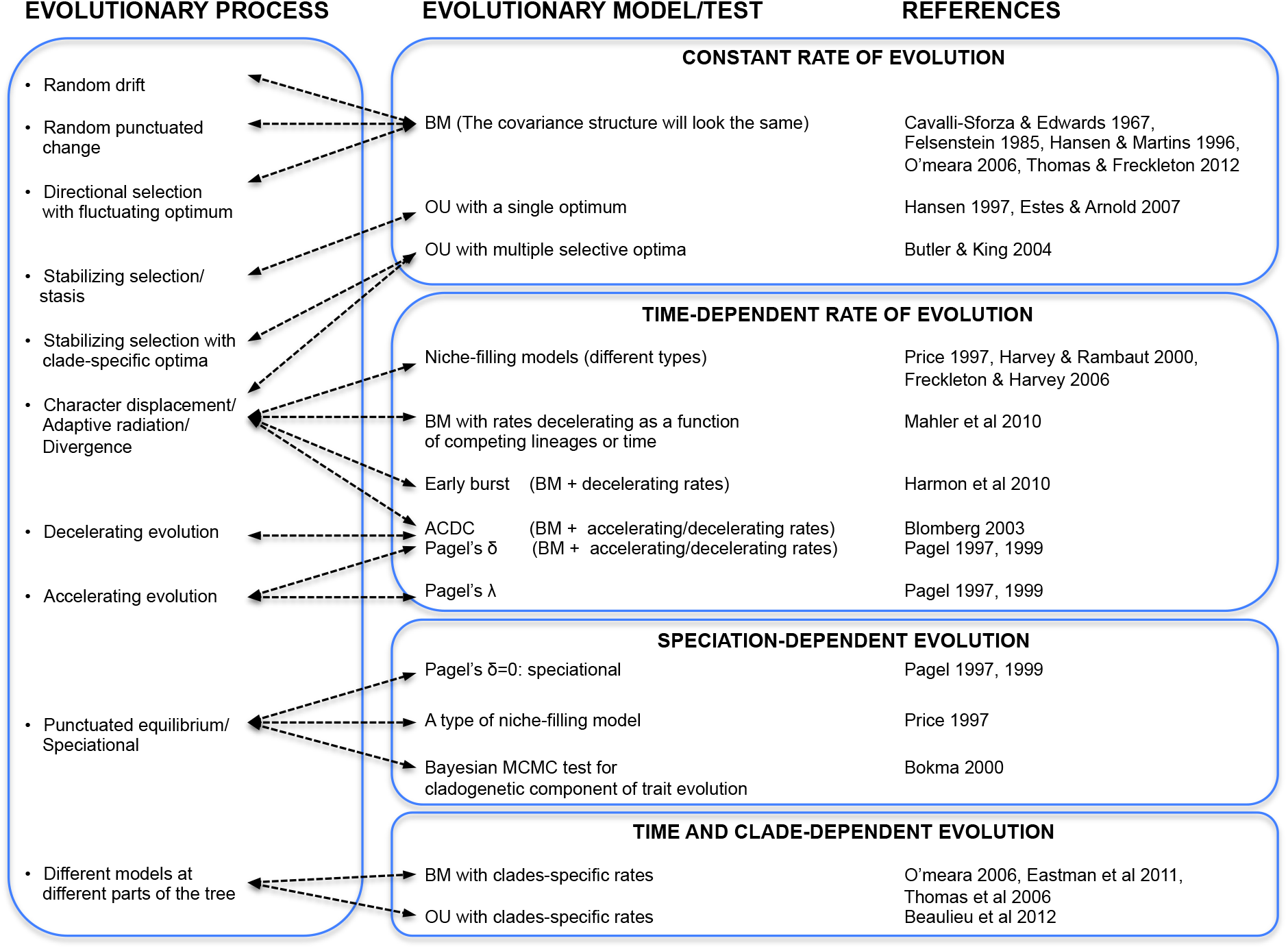
Evolutionary processes shaping the evolution of species traits are approximated by evolutionary models. Some processes, such as adaptive radiation or speciational processes can be approximated or tested by a number of slightly different models. See Box 1 for abbreviations and details.

**Figure 5:**
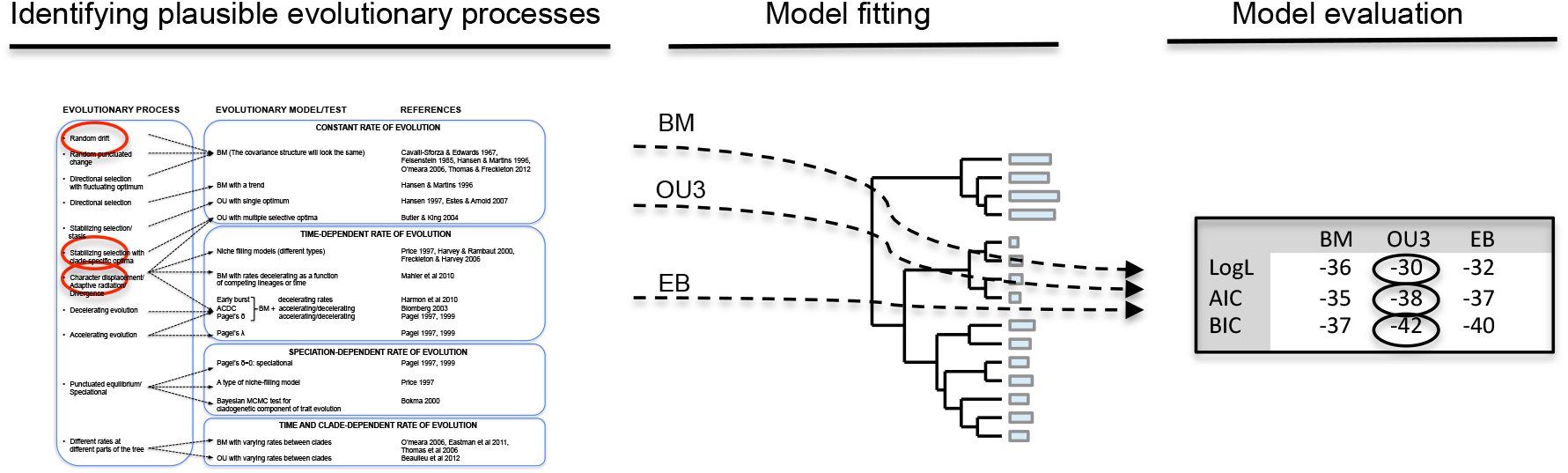
Fitting the evolutionary models to data. Based on prior biological knowledge plausible evolutionary processes responsible for shaping the evolution of species traits are identified, and the corresponding evolutionary models are fitted to the data (see Fig. 4 for a readable version of this step). The model that fits best to the data (trait values and phylogeny) is commonly selected with likelihood ratio test (LRT), or with model selection criteria such as Akaike information criterion (AIC) or Schwarz criterion (SC=BIC).

Nonetheless, recent criticism about information theoretic approaches cast a doubt on their ability to discern the correct model. As Boettiger et al. (2012) and Slater and Pennell (2013) argue, in comparative studies *predictive approaches* are more robust and powerful means for model selection. While information theoretic approaches select the model which maximizes the posterior probability of the observed values, predictive approaches prefer the model which best predicts the observed values through simulation (Slater and Pennell, 2013). In this approach the models are first fitted and parameters for each model of evolution are estimated, and subsequently used to simulate new data. The models are then evaluated based on how closely they predicted the observed data. This procedure is available in the R-package “geiger”(Harmon et al., 2008), so far to test for early burst, Brownian motion and rate shift models. Package “pmc” (Boettiger et al., 2012) allows a simulation based method to choose between models fitted in“geiger”, “ape” (Paradis et al., 2004) and “ouch” (King and Butler, 2009). Given that complex models have a higher number of parameters, the phylogenies have to be large enough to allow a reasonable inference of the evolutionary model (Boettiger et al., 2012). Thanks to the increasing availability of molecular data, phylogenetic trees are growing to include many thousand species (Bininda-Emonds et al., 2007; Smith and Donoghue, 2008; Thuiller et al., 2011; Jetz et al., 2012). With such a variety of life forms it becomes reasonable, even necessary, to assume and test much more complex models of evolution than a simple Brownian motion.

###### Evolutionary models for continuous characters

The simplest evolutionary model is the **Brownian motion model** (BM, Cavalli-Sforza and Edwards, 1967; Felsenstein, 1985, 1988). Under this model the traits are evolving randomly in any direction from the mean at each instant of time, with a net change of zero. The probability of character change is thus proportional to branch length, and the correlation among trait values at the tips of the tree decreases linearly with increasing phylogenetic distance between species (i.e. the more closely related the species, the more similar their traits are, Hansen and Martins, 1996). Exactly the same correlation structure is also expected when traits evolve under some other processes, such as directional or stabilizing selection with fluctuating optimum or punctuated change (periods of stasis alternated by abrupt changes, Hansen and Martins, 1996; O’Meara et al., 2006; Thomas and Freckleton, 2011). The assumptions of BM are violated if the values of a trait are near their biological limits and therefore cannot decrease or increase independently of the current value, or if the trait is under stabilizing selection (O’Meara et al., 2006). In those cases trait evolution is better described by the **Ornstein-Uhlenbeck model** (OU, Hansen, 1997; Butler and King, 2004; Estes and Arnold, 2007), an extension of BM which has an additional term describing the “pull” towards an optimum value (known as mean-reversion rate in finanical mathematics). When the value of this constraint equals zero, the model is equal to BM. On the other hand, the higher the pull towards an optimal value is, the lower the correlation among closely related species will be, as all species evolve towards the same optimum.

Another process of great evolutionary importance is “adaptive radiation”, which traces back to Simpson (1944). According to this process, species traits initially evolve rapidly and then slow down as the niche space becomes filled, which is basically opposing the idea of gradual evolution as described by simple BM (Harmon et al., 2003, 2010; Slater et al., 2010). It is modeled as BM with decelerating rates of evolution through time, a model commonly known as **early-burst (EB)** or **ACDC** (accelerating versus decelerating rates of character evolution, Blomberg et al., 2003), which can also be tested with **Pagel’s δ** (Pagel, 1997; Pagel et al., 1999). Another way of detecting the pattern of decelerating evolutionary rates is to infer rate shifts through phylogeny, as described in Eastman et al. (2011), or by calculating the morphological disparity index (MDI, Harmon et al., 2003).

A further option is to perform the node height test: EB occurs when the standardized independent contrasts of trait values are higher deeper in the tree than among more recent nodes in the phylogeny (Freckleton and Harvey, 2006). Although the model of adaptive radiation is generally well supported in paleontology, it has not been often observed in comparative studies (Harmon et al., 2010). Slater and Pennell (2013) argue the inability of detecting EB may be because of lack of power of currently employed methods, rather than the absence of such pattern in nature. Several other models describing adaptive radiation exist which assume a decrease of evolutionary rates as a function of the number of competing lineages (Mahler et al., 2010), or are refinements of Price’s **niche-filling** models (Price, 1997; Harvey and Rambaut, 2000; Freckleton and Harvey, 2006). Adaptive radiation can also be fitted with a stabilizing selection model where different clades in the tree evolve towards different optima (multiple-optimum OU model Butler and King, 2004).

Similarly, to investigate the tempo of evolution – whether traits evolved rapidly immediately after speciation events followed by a long period of stasis – it is necessary to fit **punctuational or speciational** models of evolution, where the evolutionary change is a function of speciation events and is independent of branch lengths (Gould and Eldredge, 1972; Huey and Bennett, 1987; Pagel, 1997; Pagel et al., 1999; Pagel, 2002). This type of evolution can also be detected by testing for a cladogenetic component of trait evolution with Bayesian MCMC test (Bokma, 2008). Models can also assign different rates of trait evolution to different parts of a tree (O’Meara et al., 2006; Thomas et al., 2006; Eastman et al., 2011; Venditti et al., 2011; Beaulieu et al., 2012; Revell, 2012). It is possible to identify the location of a rate shift in the phylogeny with R-packages “phytools” (evol.rate.mcmc Revell, 2012), or “geiger” (rjmcmc.bm).

Different evolutionary models for continuously varying trait can be fitted with R-package “geiger” (Harmon et al., 2008), “ouch” (Butler and King, 2004), ‘ouwie” (Beaulieu et al., 2012), whereas Mahler et al. (2010) model can be fitted by fitDiversityModel in phytools.

###### Discrete characters

Statistical models describing the evolution of discrete characters are based on continuous-time Markov process, equivalent to the Brownian motion model for continuous characters (Schluter et al., 1997; Cunningham et al., 1998; Pagel et al., 1999; Ronquist, 2004). The earliest and simplest such model is the Jukes-Cantor model proposed for nucleotide substitution with equal transition rates (Jukes and Cantor, 1969; Galtier et al., 2005).

Kimura (1980) extended it to a two-rate model to allow the transition rates between nucleotides to differ. Today this family of models are known as Mk models, Markov models which can assume *k* states. (Lewis, 2001). The central feature of the model is the rate matrix, which contains the instantaneous transition rates between different character states (Pagel, 1997; Pagel et al., 1999). With a 3-states trait, this transition matrix is a 3 × 3 matrix with forward and backward transition rates represented on the off-diagonals (Revell, 2014). The model can assume different rates among characters (the rate A*→* B may differ from B *→* C), and the direction of change (the forward transition A *→* B may differ from the backward direction B *→* A). Until recently, the transition rates were fixed and applied to the entire phylogeny, without the possibility, for instance, to assume a different rate of A *→* B transition on different parts of the tree. This is now possible with the R package “corHHM”, which handles the different rate classes as hidden character states (e.g. fast and slow; Beaulieu et al., 2013). Another innovative way to model the evolution of discrete characters is through the “threshold” model, first described by (Wright, 1934), in which the discrete trait is practically transformed to a continuous character, an unobserved trait called “liability” with fixed thresholds (Felsenstein, 2012; Revell, 2014). For example, a trait with two states (A, B) can be represented by a continuous scale liability axis with an arbitrary threshold (e.g. at 0), so that when the liability assumes negative values, the trait is in state A, otherwise in state B. The threshold model is biologically reasonable because it models a discrete character as a continuous trait where the probability of the character to change states decreases with time: the longer the time after the character crossed the threshold and moved to another state, the less probable it is the return to previous state, in contrast to Mk model, in which the amount of time at a certain state does not influence the probability of change (Felsenstein, 2002). The threshold model is implemented in R package “phytools” (Revell, 2012) and permits ancestral reconstruction of discrete characters under BM and OU models of evolution. Several other R packages are available for reconstruction of discrete traits under both joint and marginal methods, allowing multistate characters and different transition rates: diversitree (FitzJohn, 2012), geiger (Harmon et al., 2008), ape (Paradis et al., 2004), as well as other software such as MESQUITE, BayesTraits and SIMMAP (Bollback, 2006). J. Felsenstein’s webpage (http://evolution.genetics.washington.edu/phylip/software.html) provides an overview of other available phylogeny software.

##### Integrating fossil records

Incorporating known ancestral values dramatically improves the inference of the evolutionary model, in particular by allowing the detection of directional evolutionary trends, which are virtually unobservable with extant taxa only (Oakley and Cunningham, 2000; Finarelli and Flynn, 2006; Albert et al., 2009; Slater et al., 2012). Substantial improvements were shown for model detection for all tested models (BM, AC/DC and OU: Slater et al., 2012). Integrating prior information by directly assigning values to specific ancestral nodes in the phylogeny is now possible and technically straightforward (e.g. in R-packages “phytools” and “geiger”; Slater et al., 2012). Constraining values from wandering too far from the optimum value can be achieved by simulating “bounded evolution” by varying the “pull” parameter of the OU model, which in turn determines the width around the optimum, or by setting absolute limits (as proposed by Revell, 2007; Revell et al., 2008). Given that the interpretation of results in comparative methods largely depend on the model of evolution, it will be important to integrate all available prior knowledge from paleosciences, continue developing realistic evolutionary models, as well as establishing reliable techniques to choose among them.

#### Ancestral niche reconstruction

Rapidly developing statistical reconstruction methods permit the estimation of ancestral trait values based on its present-day value, the phylogenetic relationship among species and an evolutionary model. Therefore the inference of the best evolutionary model should be an integral part of the reconstruction procedure. If a model is not specified, most reconstruction methods will follow a BM model by default, and their output will be identical or very similar (see Table 2).

**Table 1:** Reconstruction outcome from different methods for continuous characters. Under default conditions untransformed ultrametric tree and/or no model specification - all methods will produce roughly the same ancestral states, as all assume Brownian motion model of evolution. Note: independent contrast (IC) method will yield the same ancestral state estimates as the other methods only when each node of the tree is separately re-rooted during the reconstruction process (Maddison, 1991; Garland et al., 1997). Weighted squared-change parsimony (WSqCP) and independent contrasts can also assume different models of evolution by reconstructing the trait values on a transformed phylogeny. Bayesian estimate can lead to a different result under the same model due to different prior distributions for model parameters.

|  |  |  |  |  |  |  |  |  |
| --- | --- | --- | --- | --- | --- | --- | --- | --- |
| IC [untransformed tree] | ≈ | WSqCP [untransformed tree] | = | ML [BM] | = | GLS [BM] | ≈ | Bayesian [BM] |
| IC [OU tree] | ≈ | WSqCP [OU tree] | = | ML [OU] | = | GLS [OU] | ≈ | Bayesian [OU] |

**Table 2:** A proposal of best practice and data format for phylogenetic comparative studies

|  | Best choice | Alternative |
| --- | --- | --- |
| **Phylogeny** | Sample of trees<br>Chronogram | Single tree<br>Phylogram/topology |
| **Climate niche from occurrence data** | Modelled data | Raw data |
| **Climate niche from experimental data** | Direct measures | - |
| **Niche statistics** | Empirical distribution | Point values |
| **Evolutionary model** | Selected according to prior knowledge,<br>followed by model selection based on fit | No selection |
| **Ancestral niche projection** | To paleoclimate maps, if reliable | To present-day maps |
| **Validation** | Fossil records | - |

As in model-fitting, the methods for continuous and discrete traits are slightly different. Climate variables are most commonly expressed on a **continuous** scale and ancestral climate niches can be reconstructed following the methods for continuous characters. Squared-change parsimony (or weighted squared-change parsimony) was initially the most widely used method. The optimal values for ancestral characters are found when the sum of their squared changes over the whole phylogenetic tree reaches the minimum value (Maddison, 1991; Garland et al., 1997). Weighted squared-change parsimony takes into account branch lengths (i.e. evolutionary time), so that the resulting reconstruction corresponds to BM evolution (Maddison, 1991; Webster and Purvis, 2002). Another widely used method was Felsenstein’s (1985) independent contrasts (IC). Although weighted squared-change parsimony and IC both implicitly assume a BM model of evolution, those two methods will yield slightly different reconstructed values for all nodes except the basal, because independent contrasts use “local” optimization (only daughter nodes are considered to infer the value of the ancestor), as opposed to “global” optimization used in squared-change parsimony (Maddison, 1991; Garland et al., 1997; Webster and Purvis, 2002). Nowadays more commonly used methods are maximum likelihood (ML) (Schluter et al., 1997; Cunningham et al., 1998), generalized least squares (GLS) (Grafen, 1989; Martins and Hansen, 1997; Pagel et al., 1999; Martins, 1999) and Bayesian approaches (Pagel et al., 2004). Ancestral traits, and hence also climate niches can be estimated using R-packages “ape”, “phytools” and “phyloclim” (Heibl et al., 2013) and other software as MESQUITE (Maddison and Maddison, 2001), BayesTraits (Pagel and Meade, 2007) or COMPARE (Martins, 2004). Table 3. summarizes the approaches used in climatic niche reconstruction studies, indicating methodological improvements over time.

**Table 3:**
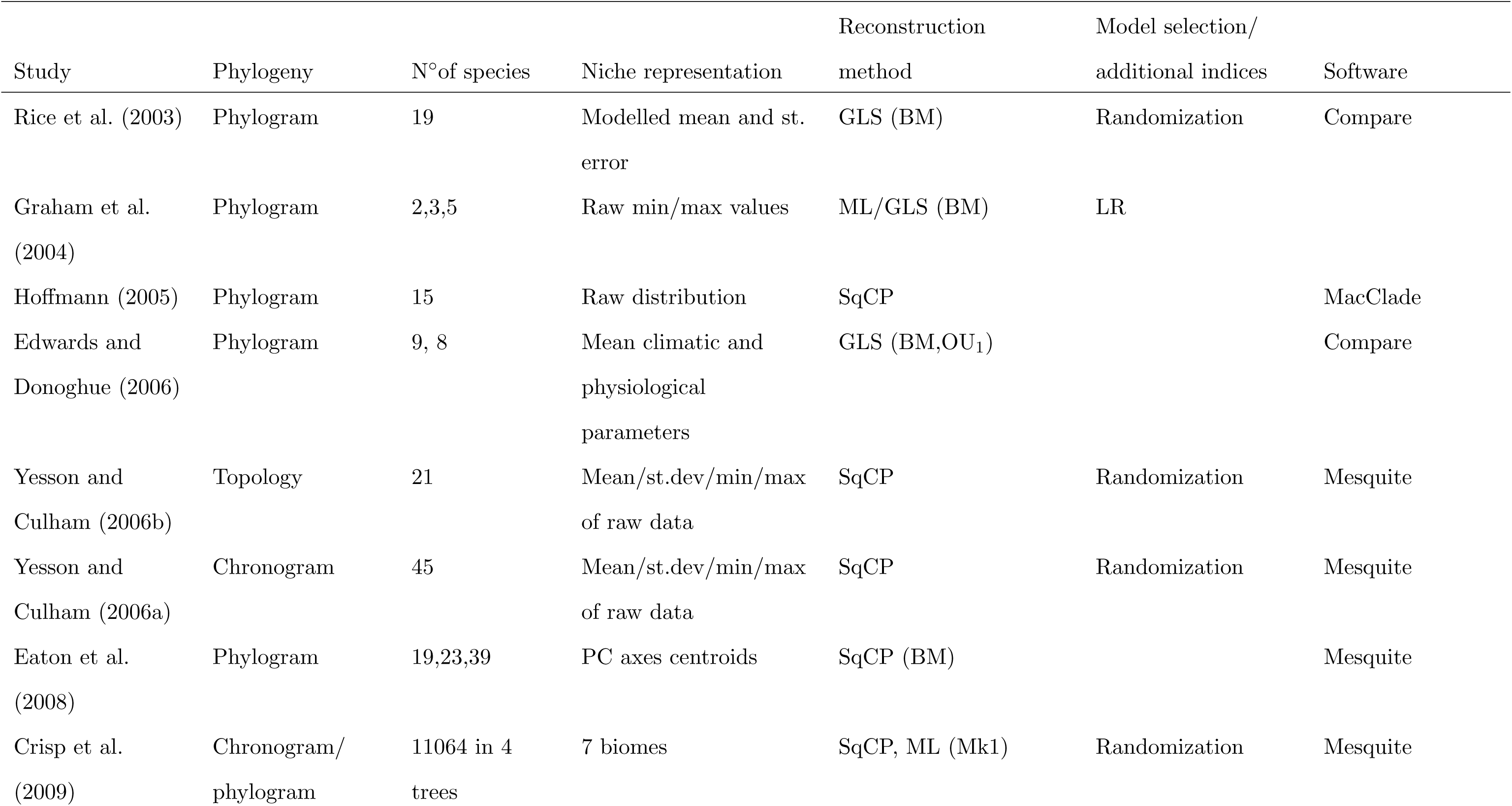
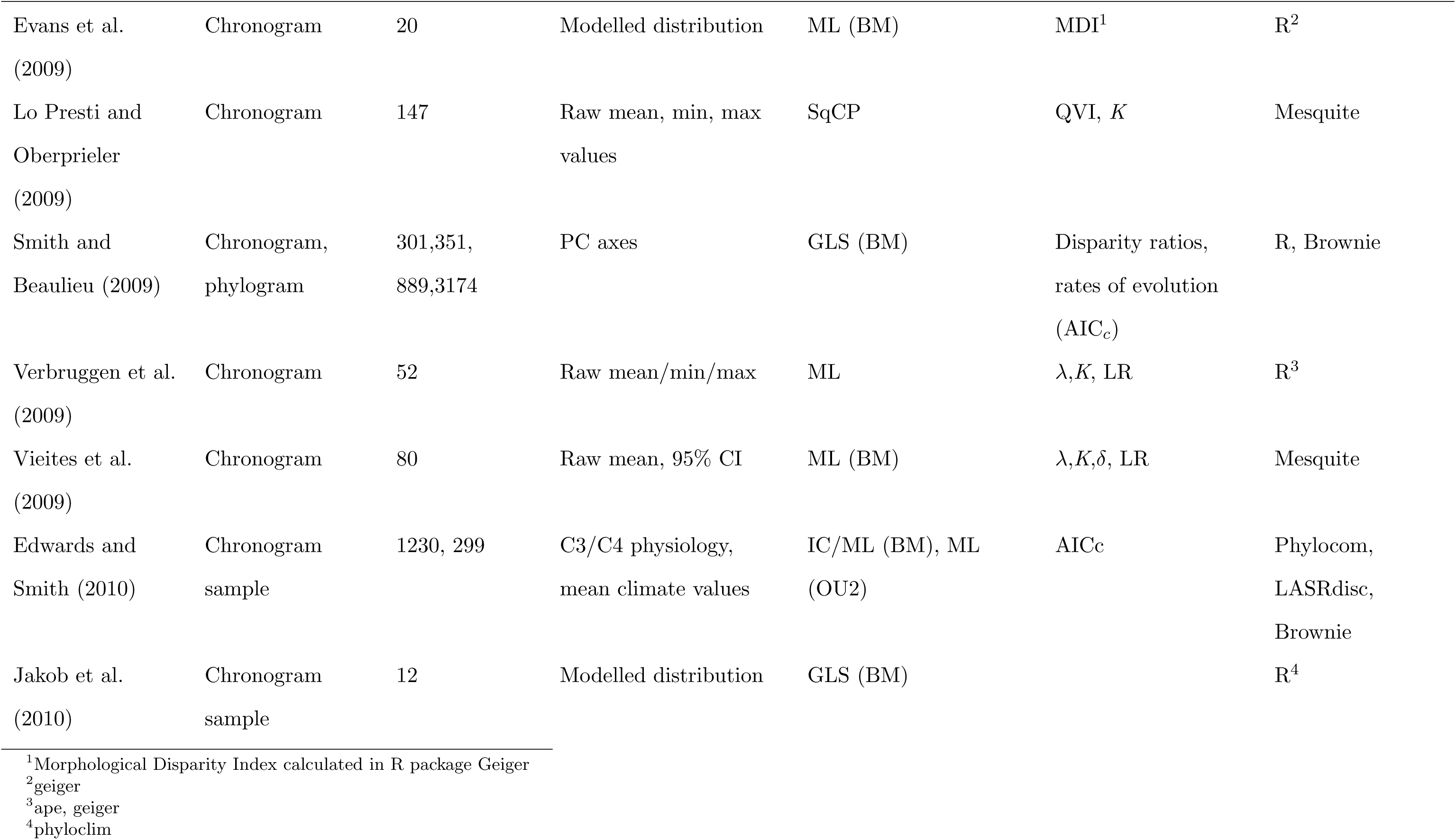
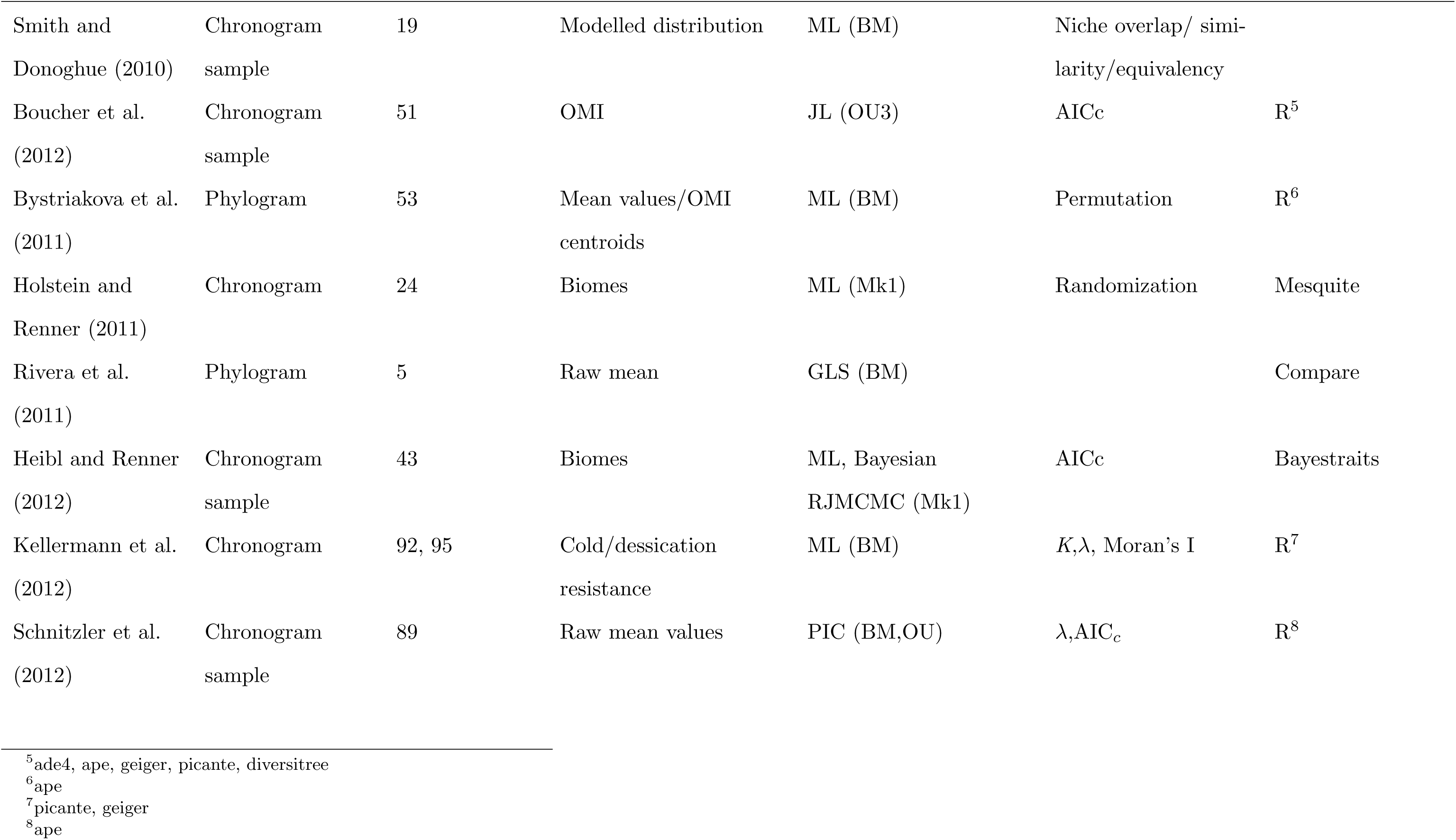
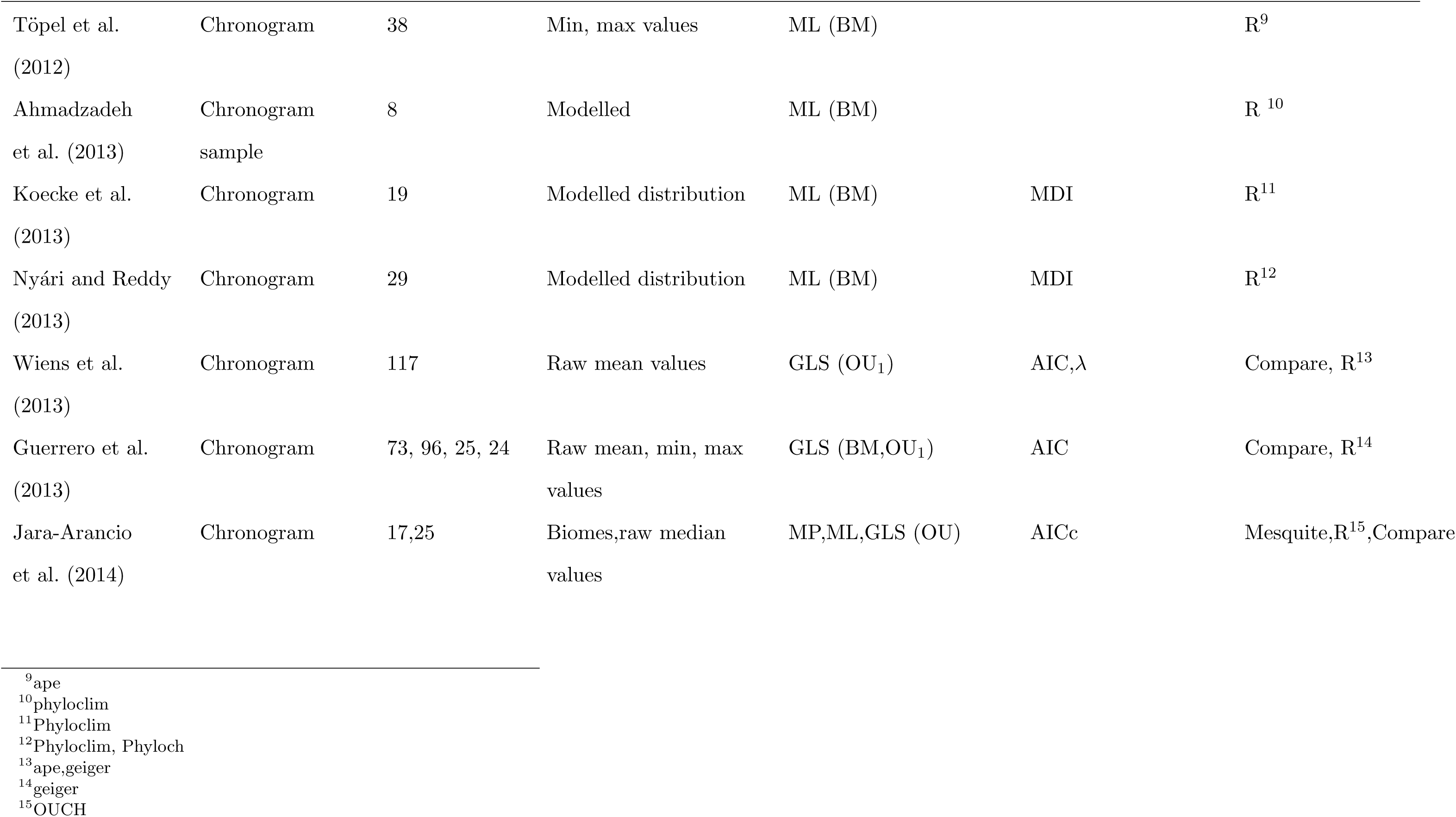
List of studies reconstructing the ancestral climate niche in chronological order. The fields with missing values indicate either that the tests were not performed or no information is available in the text. An overview of R packages implementing comparative phylogenetic methods is available in O’Meara (2012).

One could represent climate preferences as **discrete** characters (e.g. by categorising temperature values into “arid” and “mesic” conditions). Performing this categorization *after* the reconstruction of continuous climate variables allows more precision and avoids spurious results due to arbitrarily chosen thresholds. Maximum parsimony reconstructs the ancestral values by minimizing the number of changes needed to reach the observed present-day values (Pagel et al., 1999; Pedersen et al., 2007). Maximum likelihood and Bayesian methods are broadly grouped into “joint” or “global” and “marginal” or “local” reconstructions. Joint reconstruction finds the states which jointly maximize the likelihood over the whole phylogeny. In contrast, marginal reconstruction singles out the state with the highest likelihood at each node separately, which can be useful to test a specific hypothesis at a certain node in the tree (Pagel, 1999). Models which describe the evolution of discrete characters are based on the Markov-transition process of the probability of the character (see Box 1, Pagel et al., 1999).

Reconstruction procedures take into account evolutionary models by transforming the branch lengths, the path separating species pairs from their common ancestor in the phylogenetic tree. For instance, according to early burst model, evolution is faster closer to the root of the tree, therefore after transformation those branches will be longer compared to the branches closer to the tips where the rate of evolution is slowing down. Reconstructing the values according to a specific model of evolution in R can be done in two steps: first, it is necessary to transform the phylogenetic tree according to the previously tested best fitting model (e.g. rescale function in “geiger”), and afterwards this rescaled tree can be used for ML-based ancestral niche reconstruction under the default BM model of evolution (e.g. ace function in “ape”). The obtained ancestral values correspond to values evolved according to the evolutionary model used to transform the tree. Accordingly, weighted squared change parsimony or independent contrasts can fit different models of evolution in a computationally efficient way, by employing appropriate transformations of the phylogeny (see Table 1).

Regardless of the character type or method of choice, estimating trait history on a sample of possible phylogenetic trees instead of using only the single best tree allows to incorporate phylogenetic uncertainty in the analysis. Reconstructed estimates of all trees are then averaged and their distribution provides uncertainty estimates. This procedure is not limited to Bayesian analysis, but can be applied to any ancestral reconstruction method (Donoghue and Ackerly, 1996; Martins and Hansen, 1997; de Villemereuil et al., 2012).

Visualizing and validating the ancestral range and climatic niche If the aim of niche reconstruction was exploring the unfolding of evolutionary changes through time, we may want to visualize the ancestral climatic niche in an abstract multidimensional climate space (e.g. Veloz et al., 2012). On the other hand, to visualize the historical distribution and appreciate the extent of range shift through time, ancestral niche is often projected to a geographic map along with the current climatic niche (e.g. Yesson and Culham, 2006a; Lawing and Polly, 2011; Töpel et al., 2012). This may be problematic because environmental variables today are most likely correlated differently among each other than they were in the past (Boucher et al., 2012), given that non-analog climate conditions were already present at different time steps even on a short time scale, such as in the Quaternary (Jackson and Overpeck, 2000; Williams and Jackson, 2007). Therefore, projecting the ancestral niche to today’s world and vice-versa will not accurately represent the ancestral range, as parts of the range may be missing while some other areas might be wrongly assigned. It practically shows where the ancestor would live today, but not necessarily where it actually lived in the past. The ideal solution would be to project to paleoclimate maps, but because they become less reliable the further one goes back in time, it is difficult to infer the correct ancestral range solely with SDMs, without fossil records. Fossil records are scarce for most species and are often biased with respect to climate, topography, species size and abundance, yielding fewer traces of rare and small-sized animal species especially in wet tropical climates (Kidwell and Flessa, 1995). Still, whenever available they are a valuable indicator of a species’ past distribution as they are generally buried within the species’ range so spatial displacements between past and present are likely due to a range shift of species (Kidwell and Flessa, 1995). Fossil records of species occurrences can prove that species were present in the study area at the specific time (Vieites et al., 2009), and/or in the predicted ancestral range.

Known ancestral climatic values from paleodata add valuable information and should either be used to aid a better approximation of evolutionary models by constraining specific nodes to known values, as previously seen, or be employed for validating the reconstructed niche values. Validation may be more important when current data already yield a well-constrained model. In contrast, poorly defined models may profit from integrating paleoclimate data into the estimation process.

## Summary and recommendations

Here we summarize and propose tentative guidelines for optimal use of species occurrence and climatic data in phylogenetic comparative studies.

### Niche representation

- When present-day climate niches are inferred from spatial occurrences, niche models are a better choice than raw data.
- Each climatic variable should ideally be expressed by species preference for the full range of values (i.e. empirical densities), instead of being summarized by the mean or other point estimates (e.g. mean temperature).

### Ancestral niche reconstruction

- Before reconstructing the niche, a best-fitting evolutionary model should be estimated for each climatic variable.
- Known paleoclimate data can either be used to improve the evolutionary model inference or to validate the reconstructed values.
- Evolutionary changes are best visualised in an abstract climatic space.
- Ancestral species range should be projected to a paleoclimate map.

## Concluding remarks

Analysing the evolution of climatic niches integrates species distribution modeling, phylogenies, evolutionary models as well as elements of paleosciences. Such a complex research question requires careful consideration of each component to minimize potential bias and information loss. We primarily focused and discussed the most appropriate methods to represent the climatic niche through species distribution modeling and outlined the procedure for ancestral niche reconstruction. This research field has a lot to gain from improvements in other areas, particularly from developing new evolutionary models, which would better approximate processes on macroevolutionary scale. Available paleodata has the potential to greatly improve the detection of evolutionary models (Slater et al., 2012), and we expect the future efforts in this interdisciplinary field to focus especially on a better integration of phylogenetic, paleontological and climatic data.

### Acknowledgments

We would like to thank Jan Schnitzler and Marten Winter for providing valuable comments that greatly improved the manuscript. This research was supported by the German Research Foundation, DFG (DO 786/5-1). The authors declare no conflict of interest.

## References

Ackerly, D. D., Schwilk, D. W. and Webb, C. O. 2006. Niche evolution and adaptive radiation: testing the order of trait divergence. – Ecology 87: S50–S61.

Afkhami, M. E., McIntyre, P. J. and Strauss, S. Y. 2014. Mutualist-mediated effects on species’ range limits across large geographic scales. – Ecology letters 17(10): 1265–1273.

Ahmadzadeh, F., Flecks, M., Carretero, M. A., Böhme, W., Ilgaz, C., Engler, J. O., James Harris, D., Üzüm, N. and Rödder, D. 2013. Rapid lizard radiation lacking niche conservatism: ecological diversification within a complex landscape. – Journal of Biogeography 40(9): 1807–1818.

Akaike, H. 1973. Information theory and an extension of the maximum likelihood principle. – In: Second International Symposium on Information Theory. Springer Verlag, vol. 1, pp.267–281.

Albert, J. S., Johnson, D. M. and Knouft, J. H. 2009. Fossils provide better estimates of ancestral body size than do extant taxa in fishes. – Acta Zoologica 90(1): 357–384.

Beaulieu, J. M., Jhwueng, D.-C., Boettiger, C. and O’Meara, B. C. 2012. Modeling stabilizing selection: expanding the Ornstein–Uhlenbeck model of adaptive evolution. – Evolution 66(8): 2369–2383.

Beaulieu, J. M., O’Meara, B. C. and Donoghue, M. J. 2013. Identifying hidden rate changes in the evolution of a binary morphological character: the evolution of plant habit in campanulid angiosperms. – Systematic Biology 62(5): 725–737.

Bininda-Emonds, O., Cardillo, M., Jones, K., MacPhee, R., Beck, R., Grenyer, R., Price, S., Vos, R., Gittleman, J. and Purvis, A. 2007. The delayed rise of present-day mammals. – Nature 446(7135): 507–512.

Bininda-Emonds, O. R. 2004. Phylogenetic supertrees: combining information to reveal the tree of life. – vol. 4. Springer, Berlin.

Blomberg, S., Garland Jr, T. and Ives, A. 2003. Testing for phylogenetic signal in comparative data: behavioral traits are more labile. – Evolution 57(4): 717–745.

Boettiger, C., Coop, G. and Ralph, P. 2012. Is your phylogeny informative? Measuring the power of comparative methods. – Evolution 66(7): 2240–2251.

Bokma, F. 2008. Detection of “punctuated equilibrium” by Bayesian estimation of speciation and extinction rates, ancestral character states, and rates of anagenetic and cladogenetic evolution on a molecular phylogeny. – Evolution 62(11): 2718–2726.

Bollback, J. P. 2006. SIMMAP: stochastic character mapping of discrete traits on phylogenies. – BMC Bioinformatics 7(1): 88.

Boucher, F., Thuiller, W., Roquet, C., Douzet, R., Aubert, S., Alvarez, N. and Lavergne, S. 2012. Reconstructing the origins of high-alpine niches and cushion life form in the genus *Androsace* S.L. (Primulaceae). – Evolution 66(4): 1255–1268.

Boucher, F. C., Thuiller, W., Davies, T. J. and Lavergne, S. 2014. Neutral biogeography and the evolution of climatic niches. – The American Naturalist 183(5): 573.

Bruno, J. F., Stachowicz, J. J. and Bertness, M. D. 2003. Inclusion of facilitation into ecological theory. – Trends in Ecology & Evolution 18(3): 119–125.

Burnham, K. P. and Anderson, D. R. 2002. Model selection and multi-model inference: a practical information-theoretic approach. – Springer, Berlin.

Butler, M. and King, A. 2004. Phylogenetic comparative analysis: a modeling approach for adaptive evolution. – The American Naturalist 164(6): 683–695.

Bystriakova, N., Schneider, H. and Coomes, D. 2011. Evolution of the climatic niche in scaly tree ferns (Cyatheaceae, Polypodiopsida). – Botanical Journal of the Linnean Society 165(1): 1–19.

Cavalli-Sforza, L. L. and Edwards, A. W. 1967. Phylogenetic analysis. Models and estimation procedures. – American Journal of Human Genetics 19(3 Pt 1): 233.

Cooper, N., Jetz, W. and Freckleton, R. 2010. Phylogenetic comparative approaches for studying niche conservatism. – Journal of Evolutionary Biology 23(12): 2529–2539.

Crisp, M., Arroyo, M., Cook, L., Gandolfo, M., Jordan, G., McGlone, M., Weston, P., Westoby, M., Wilf, P. and Linder, H. 2009. Phylogenetic biome conservatism on a global scale. – Nature 458(7239): 754–756.

Crisp, M. D. and Cook, L. G. 2012. Phylogenetic niche conservatism: what are the underlying evolutionary and ecological causes?. – New Phytologist 196(3): 681–694.

Cunningham, C., Omland, K. and Oakley, T. 1998. Reconstructing ancestral character states: a critical reappraisal. – Trends in Ecology & Evolution 13(9): 361–366.

Donoghue, M. J. and Ackerly, D. D. 1996. Phylogenetic uncertainties and sensitivity analyses in comparative biology. – Philosophical Transactions of the Royal Society of London. Series B 351(1345): 1241–1249.

Dormann, C. F., Gruber, B., Winter, M. and Herrmann, D. 2010. Evolution of climate niches in European mammals?. – Biology Letters 6(2): 229–232.

Eastman, J. M., Alfaro, M. E., Joyce, P., Hipp, A. L. and Harmon, L. J. 2011. A novel comparative method for identifying shifts in the rate of character evolution on trees. – Evolution 65(12): 3578–3589.

Eaton, M., Soberón, J. and Peterson, A. 2008. Phylogenetic perspective on ecological niche evolution in American blackbirds (family Icteridae). – Biological Journal of the Linnean Society 94(4): 869–878.

Edwards, E. and Donoghue, M. 2006. Pereskia and the origin of the cactus life-form. – The American Naturalist 167(6): 777–793.

Edwards, E. and Smith, S. 2010. Phylogenetic analyses reveal the shady history of C4 grasses. – Proceedings of the National Academy of Sciences, USA 107(6): 2532–2537.

Elith, J., Graham*, C., Anderson, R., Dudik, M., Ferrier, S., Guisan, A., Hijmans, R., Huettmann, F., Leathwick, J., Lehmann, A. et al. 2006. Novel methods improve prediction of species’ distributions from occurrence data. – Ecography 29(2): 129–151.

Estes, S. and Arnold, S. J. 2007. Resolving the paradox of stasis: models with stabilizing selection explain evolutionary divergence on all timescales. – The American Naturalist 169(2): 227–244.

Evans, M., Hearn, D., Hahn, W., Spangle, J. and Venable, D. 2005. Climate and life-history evolution in evening primroses (*Oenothera*, Onagraceae): a phylogenetic comparative analysis. – Evolution 59(9): 1914–1927.

Evans, M., Smith, S., Flynn, R. and Donoghue, M. 2009. Climate, niche evolution, and diversification of the “bird-cage” evening primroses (*Oenothera*, Sections *Anogra* and *Kleinia*). – The American Naturalist 173(2): 225–240.

Felsenstein, J. 1985. Phylogenies and the comparative method. – American Naturalist 125(1): 1–15.

Felsenstein, J. 1988. Phylogenies and quantitative characters. – Annual Review of Ecology and Systematics 19: 445–471.

Felsenstein, J. 2002. Quantitative characters, phylogenies, and morphometrics. – In: MacLeod, N. and Forey, P. L. (eds.), Morphology, shape and phylogeny. Taylor and Francis, London and New York, pp. 27–44.

Felsenstein, J. 2005. Using the quantitative genetic threshold model for inferences between and within species. – Philosophical Transactions of the Royal Society B 360(1459): 1427–1434.

Felsenstein, J. 2012. A comparative method for both discrete and continuous characters using the threshold model. – The American Naturalist 179(2): 145–156.

Finarelli, J. A. and Flynn, J. J. 2006. Ancestral state reconstruction of body size in the Caniformia (Carnivora, Mammalia): the effects of incorporating data from the fossil record. – Systematic Biology 55(2): 301–313.

FitzJohn, R. G. 2012. Diversitree: comparative phylogenetic analyses of diversification in R. – Methods in Ecology and Evolution 3(6): 1084–1092.

Franklin, J. and Miller, J. 2009. Mapping species distributions: spatial inference and prediction. – vol. 338. Cambridge University Press Cambridge.

Freckleton, R. P. and Harvey, P. H. 2006. Detecting non-Brownian trait evolution in adaptive radiations. – PLoS biology 4(11): e373.

Galtier, N., Gascuel, O. and Jean-Marie, A. 2005. Markov models in molecular evolution. – In: Nielsen, R. (ed.), Statistical Methods in Molecular Evolution. Springer, New York, pp. 3–24.

Garland, T., Martin, K. and Diaz-Uriarte, R. 1997. Reconstructing ancestral trait values using squared-change parsimony: plasma osmolarity at the origin of amniotes. – In: Sumida, S. & and Martin, K. L. (eds.), Amniote Origins: Completing the Transition to Land. Academic Press San Diego, CA, pp. 425–501.

Gould, S. J. and Eldredge, N. 1972. Punctuated equilibria: an alternative to phyletic gradualism. – In: Schopf, T. J. (ed.), Models in paleobiology. Freeman, Cooper, San Francisco, pp. 82–115.

Grafen, A. 1989. The phylogenetic regression. – Philosophical Transactions of the Royal Society of London. Series B 326: 119–157.

Graham, C., Ron, S., Santos, J., Schneider, C. and Moritz, C. 2004. Integrating phylogenetics and environmental niche models to explore speciation mechanisms in dendrobatid frogs. – Evolution 58(8): 1781–1793.

Guerrero, P. C., Rosas, M., Arroyo, M. T. and Wiens, J. J. 2013. Evolutionary lag times and recent origin of the biota of an ancient desert (Atacama–Sechura). – Proceedings of the National Academy of Sciences 110(28): 11469–11474.

Guisan, A. and Zimmermann, N. 2000. Predictive habitat distribution models in ecology. – Ecological modelling 135(2): 147–186.

Hansen, T. 1997. Stabilizing selection and the comparative analysis of adaptation. – Evolution 51(5): 1341–1351.

Hansen, T. and Martins, E. 1996. Translating between microevolutionary process and macroevolutionary patterns: the correlation structure of interspecific data. – Evolution 50: 1404–1417.

Hardy, C. 2006. Reconstructing ancestral ecologies: challenges and possible solutions. – Diversity and Distributions 12(1): 7–19.

Hardy, C. and Linder, H. 2005. Intraspecific variability and timing in ancestral ecology reconstruction: a test case from the Cape flora. – Systematic Biology 54(2): 299–316.

Harmon, L., Schulte II, J., Larson, A. and Losos, J. 2003. Tempo and mode of evolutionary radiation in iguanian lizards. – Science 301(5635): 961–964.

Harmon, L. J., Losos, J. B., Jonathan Davies, T., Gillespie, R. G., Gittleman, J. L., Bryan Jennings, W., Kozak, K. H., McPeek, M. A., Moreno-Roark, F., Near, T. J. et al. 2010. Early bursts of body size and shape evolution are rare in comparative data. – Evolution 64(8): 2385–2396.

Harmon, L. J., Weir, J. T., Brock, C. D., Glor, R. E. and Challenger, W. 2008. GEIGER: investigating evolutionary radiations. – Bioinformatics 24(1): 129–131.

Harvey, P. H. and Rambaut, A. 2000. Comparative analyses for adaptive radiations. – Philosophical Transactions of the Royal Society of London. Series B 355(1403): 1599–1605.

Heibl, C., Calenge, C. and Heibl, M. C. 2013. Package ?phyloclim?. –.

Heibl, C. and Renner, S. S. 2012. Distribution models and a dated phylogeny for Chilean *Oxalis* species reveal occupation of new habitats by different lineages, not rapid adaptive radiation. – Systematic Biology 61(5): 823–834.

Hoffmann, M. 2005. Evolution of the realized climatic niche in the genus: *Arabidopsis*(Brassicaceae). – Evolution 59(7): 1425–1436.

Holder, M., Lewis, P. et al. 2003. Phylogeny estimation: traditional and Bayesian approaches. – Nature Reviews Genetics 4(4): 275–284.

Holstein, N. and Renner, S. 2011. A dated phylogeny and collection records reveal repeated biome shifts in the African genus *Coccinia* (Cucurbitaceae). – BMC Evolutionary Biology 11(1): 28.

Huey, R. B. and Bennett, A. F. 1987. Phylogenetic studies of coadaptation: preferred temperatures versus optimal performance temperatures of lizards. – Evolution 41: 1098–1115.

Hurlbert, S. 1978. The measurement of niche overlap and some relatives. – Ecology 59(1): 67–77.

Hutchinson, G. 1957. Cold Spring Harbor Symposium on Quantitative Biology. – Concluding remarks, Quant. Biol. 22: 415–427.

Hutchinson, G. E. 1978. An Introduction to Population Ecology. – vol. 260. Yale University Press, New Haven.

Jackson, S. T. and Overpeck, J. T. 2000. Responses of plant populations and communities to environmental changes of the late Quaternary. – Paleobiology 26(4): 194–220.

Jakob, S. S., Heibl, C., Rödder, D. and Blattner, F. R. 2010. Population demography influences climatic niche evolution: evidence from diploid American *Hordeum* species (Poaceae). – Molecular Ecology 19(7): 1423–1438.

Jara-Arancio, P., Arroyo, M. T., Guerrero, P. C., Hinojosa, L. F., Arancio, G. and Méndez, M.A. 2014. Phylogenetic perspectives on biome shifts in *Leucocoryne* (Alliaceae) in relation to climatic niche evolution in western South America. – Journal of Biogeography 41(2): 328–338.

Jetz, W., Thomas, G., Joy, J., Hartmann, K. and Mooers, A. 2012. The global diversity of birds in space and time. – Nature 491(7424): 444–448.

Johnson, J. and Omland, K. 2004. Model selection in ecology and evolution. – Trends in Ecology & Evolution 19(2): 101–108.

Jukes, T. H. and Cantor, C. R. 1969. Evolution of protein molecules. – In: Munro, H. N. (ed.), Manmmalian Protein Metabolism. Academic Press, New York, pp. 21–132.

Kearney, M. and Porter, W. P. 2004. Mapping the fundamental niche: physiology, climate, and the distribution of a nocturnal lizard. – Ecology 85(11): 3119–3131.

Kellermann, V., Loeschcke, V., Hoffmann, A., Kristensen, T., Fløjgaard, C., David, J., Svenning, J. and Overgaard, J. 2012. Phylogenetic constraints in key functional traits behind species’ climate niches: patterns of desiccation and cold resistance across 95 *Drosophila* species. – Evolution 66(11): 3377–3389.

Kidwell, S. M. and Flessa, K. W. 1995. The quality of the fossil record: populations, species, and communities. – Annual Review of Ecology and Systematics 26: 269–299.

Kimura, M. 1980. A simple method for estimating evolutionary rates of base substitutions through comparative studies of nucleotide sequences. – Journal of Molecular Evolution 16(2): 111–120.

King, A. and Butler, M. 2009. ouch: Ornstein-Uhlenbeck models for phylogenetic comparative hypotheses (R package). –.

Koecke, A. V., Muellner-Riehl, A. N., Pennington, T. D., Schorr, G. and Schnitzler, J. 2013. Niche evolution through time and across continents: The story of Neotropical *Cedrela*(Meliaceae). – American Journal of Botany 100(9): 1800–1810.

Kozak, K., Graham, C. and Wiens, J. 2008. Integrating GIS-based environmental data into evolutionary biology. – Trends in Ecology & Evolution 23(3): 141–148.

Kozak, K. and Wiens, J. 2010a. Accelerated rates of climatic-niche evolution underlie rapid species diversification. – Ecology Letters 13(11): 1378–1389.

Kozak, K. H. and Wiens, J. J. 2010b. Niche conservatism drives elevational diversity patterns in Appalachian salamanders.. – The American Naturalist 176(1): 40–54.

Kraft, N., Cornwell, W., Webb, C. and Ackerly, D. 2007. Trait evolution, community assembly, and the phylogenetic structure of ecological communities. – The American Naturalist 170(2): 271–283.

Lawing, A. and Polly, P. 2011. Pleistocene Climate, Phylogeny, and Climate Envelope Models: An Integrative Approach to Better Understand Species’ Response to Climate Change. – PLoS one 6(12): e28554.

Lewis, P. O. 2001. A likelihood approach to estimating phylogeny from discrete morphological character data. – Systematic Biology 50(6): 913–925.

Lo Presti, R. M. and Oberprieler, C. 2009. Evolutionary history, biogeography and eco-climatological differentiation of the genus Anthemis L.(Compositae, Anthemideae) in the circum-Mediterranean area. – Journal of Biogeography 36(7): 1313–1332.

Losos, J. 2008a. Phylogenetic niche conservatism, phylogenetic signal and the relationship between phylogenetic relatedness and ecological similarity among species. – Ecology Letters 11(10): 995–1003.

Losos, J. B. 2008b. Rejoinder to Wiens (2008): phylogenetic niche conservatism, its occurrence and importance. – Ecology Letters 11(10): 1005–1007.

Losos, J. B. 2011. Seeing the forest for the trees: the limitations of phylogenies in comparative biology. – The American Naturalist 177(6): 709–727.

Maddison, W. 1991. Squared-change parsimony reconstructions of ancestral states for continuous-valued characters on a phylogenetic tree. – Systematic Biology 40(3): 304–314.

Maddison, W. P. and Maddison, D. R. 2001. Mesquite: a modular system for evolutionary analysis.. –.

Mahler, D. L., Revell, L. J., Glor, R. E. and Losos, J. B. 2010. Ecological opportunity and the rate of morphological evolution in the diversification of Greater Antillean anoles. – Evolution 64(9): 2731–2745.

Manly, B. F., McDonald, L. L. and Thomas, D. L. 1993. Resource selection by animals: Statistical design and analysis for field studies. – Chapman & Hall (London and New York).

Martins, E. 1999. Estimation of ancestral states of continuous characters: a computer simulation study. – Systematic Biology 48(3): 642–650.

Martins, E. 2004. COMPARE, version 4.6 b. – Computer programs for the statistical analysis of comparative data. Distributed by the author at http://compare.bio.indiana.edu/. Department of Biology, Indiana University, Bloomington, IN.

Martins, E. and Hansen, T. 1997. Phylogenies and the comparative method: a general approach to incorporating phylogenetic information into the analysis of interspecific data. – American Naturalist 149(4): 646–667.

Matthiopoulos, J., Hebblewhite, M., Aarts, G. and Fieberg, J. 2011. Generalized functional responses for species distributions. – Ecology 92(3): 583–589.

Nyári, Á. S. and Reddy, S. 2013. Comparative Phyloclimatic Analysis and Evolution of Ecological Niches in the Scimitar Babblers (Aves: Timaliidae: Pomatorhinus). – PloS one 8(2): e55629.

Oakley, T. H. and Cunningham, C. W. 2000. Independent contrasts succeed where ancestor reconstruction fails in a known bacteriophage phylogeny. – Evolution 54(2): 397–405.

O’Meara, B., Ané, C., Sanderson, M. and Wainwright, P. 2006. Testing for different rates of continuous trait evolution using likelihood. – Evolution 60(5): 922–933.

O’Meara, B. C. 2012. CRAN task view: phylogenetics. Version 2014-07-17. http://cran.r-project.org/web/views/Phylogenetics.html. –.

Pagel, M. 1997. Inferring evolutionary processes from phylogenies. – Zoologica Scripta 26(4): 331–348.

Pagel, M. 1999. The maximum likelihood approach to reconstructing ancestral character states of discrete characters on phylogenies. – Systematic Biology 48(3): 612–622.

Pagel, M. 2002. Modelling the evolution of continuously varying characters on phylogenetic trees: the case of Hominid cranial capacity. – In: MacLeod, N. and Forey, P. L. (eds.), Morphology, shape and phylogeny. Taylor and Francis, London and New York, pp. 269–286.

Pagel, M. and Meade, A. 2007. BayesTraits. – Computer program and documentation available at http://www.evolution.rdg.ac.uk/BayesTraits.html.

Pagel, M., Meade, A. and Barker, D. 2004. Bayesian estimation of ancestral character states on phylogenies. – Systematic Biology 53(5): 673–684.

Pagel, M. et al. 1999. Inferring the historical patterns of biological evolution. – Nature 401(6756): 877–884.

Paradis, E., Claude, J. and Strimmer, K. 2004. APE: analyses of phylogenetics and evolution in R language. – Bioinformatics 20: 289–290.

Pearman, P. B., D’Amen, M., Graham, C. H., Thuiller, W. and Zimmermann, N. E. 2010. Within-taxon niche structure: niche conservatism, divergence and predicted effects of climate change. – Ecography 33(6): 990–1003.

Pedersen, N., Holyoak, D. and Newton, A. 2007. Systematics and morphological evolution within the moss family Bryaceae: A comparison between parsimony and Bayesian methods for reconstruction of ancestral character states. – Molecular Phylogenetics and Evolution 43(3): 891–907.

Pennell, M. W. and Harmon, L. J. 2013. An integrative view of phylogenetic comparative methods: connections to population genetics, community ecology, and paleobiology. – Annals of the New York Academy of Sciences 1289(1): 90–105.

Peterson, A. T., Soberón, J., Pearson, R. G., Anderson, R. P., Martínez-Meyer, E., Nakamura, and Araújo, M. B. 2011. Ecological niches and geographic distributions. – Princeton University Press, Princeton.

Posada, D. and Buckley, T. R. 2004. Model selection and model averaging in phylogenetics: advantages of Akaike information criterion and Bayesian approaches over likelihood ratio tests. – Systematic Biology 53(5): 793–808.

Price, T. 1997. Correlated evolution and independent contrasts. – Philosophical Transactions of the Royal Society of London. Series B 352(1352): 519–529.

Réale, D., McAdam, A. G., Boutin, S. and Berteaux, D. 2003. Genetic and plastic responses of a northern mammal to climate change. – Proceedings of the Royal Society of London. Series B 270(1515): 591–596.

Revell, L. 2007. Testing the genetic constraint hypothesis in a phylogenetic context: a simulation study. – Evolution 61(11): 2720–2727.

Revell, L., Harmon, L. and Collar, D. 2008. Phylogenetic signal, evolutionary process, and rate. – Systematic Biology 57(4): 591–601.

Revell, L. J. 2012. phytools: an R package for phylogenetic comparative biology (and other things). – Methods in Ecology and Evolution 3(2): 217–223.

Revell, L. J. 2014. Ancestral character estimation under the threshold model from quantitative genetics. – Evolution 68(3): 743–759.

Rice, N., Martínez-Meyer, E. and Peterson, A. 2003. Ecological niche differentiation in the Aphelocoma jays: a phylogenetic perspective. – Biological Journal of the Linnean Society 80(3): 369–383.

Rivera, P., Di Cola, V., Martínez, J., Gardenal, C. and Chiaraviglio, M. 2011. Species Delimitation in the Continental Forms of the Genus *Epicrates* (Serpentes, Boidae) Integrating Phylogenetics and Environmental Niche Models. – PLoS one 6(9): e22199.

Ronquist, F. 2004. Bayesian inference of character evolution. – Trends in Ecology & Evolution 19(9): 475–481.

Roquet, C., Thuiller, W. and Lavergne, S. 2012. Building megaphylogenies for macroecology: taking up the challenge. – Ecography 36(1): 013–026.

Schluter, D., Price, T., Mooers, A. and Ludwig, D. 1997. Likelihood of ancestor states in adaptive radiation. – Evolution 51(6): 1699–1711.

Schnitzler, J., Graham, C. H., Dormann, C. F., Schiffers, K. and Peter Linder, H. 2012. Climatic niche evolution and species diversification in the Cape flora, South Africa. – Journal of Biogeography 39(12): 2201–2211.

Schwarz, G. 1978. Estimating the dimension of a model. – The Annals of Statistics 6(2): 461–464.

Seddon, A. W., Mackay, A. W., Baker, A. G., Birks, H. J. B., Breman, E., Buck, C. E., Ellis, E. C., Froyd, C. A., Gill, J. L., Gillson, L. et al. 2014. Looking forward through the past: identification of 50 priority research questions in palaeoecology. – Journal of Ecology 102(1): 256–267.

Simpson, G. G. 1944. Tempo and mode in evolution. – vol. 15. Columbia University Press, New York.

Slater, G. J., Harmon, L. J. and Alfaro, M. E. 2012. Integrating fossils with molecular phylogenies improves inference of trait evolution. – Evolution 66(12): 3931–3944.

Slater, G. J. and Pennell, M. W. 2013. Robust Regression and Posterior Predictive Simulation Increase Power to Detect Early Bursts of Trait Evolution. – Systematic Biology p. syt066.

Slater, G. J., Price, S. A., Santini, F. and Alfaro, M. E. 2010. Diversity versus disparity and the radiation of modern cetaceans. – Proceedings of the Royal Society B: Biological Sciences 277(1697): 3097–3104.

Smith, S. and Beaulieu, J. 2009. Life history influences rates of climatic niche evolution in flowering plants. – Proceedings of the Royal Society B 276(1677): 4345–4352.

Smith, S. and Donoghue, M. 2010. Combining historical biogeography with niche modeling in the *Caprifolium* clade of *Lonicera* Caprifoliaceae, Dipsacales). – Systematic Biology 59(3): 322–341.

Smith, S. A. and Donoghue, M. J. 2008. Rates of molecular evolution are linked to life history in flowering plants. – Science 322(5898): 86–89.

Soberón, J. and Nakamura, M. 2009. Niches and distributional areas: concepts, methods, and assumptions. – Proceedings of the National Academy of Sciences 106(Supplement 2): 19644–19650.

Thomas, G. and Freckleton, R. 2011. MOTMOT: models of trait macroevolution on trees. – Methods in Ecology and Evolution 3(1): 145–151.

Thomas, G. H., Freckleton, R. P. and Székely, T. 2006. Comparative analyses of the influence of developmental mode on phenotypic diversification rates in shorebirds. – Proceedings of the Royal Society B 273(1594): 1619–1624.

Thuiller, W., Lavergne, S., Roquet, C., Boulangeat, I., Lafourcade, B. and Araujo, M. 2011. Consequences of climate change on the tree of life in Europe. – Nature 470(7335): 531–534.

Töpel, M., Antonelli, A., Yesson, C. and Eriksen, B. 2012. Past Climate Change and Plant Evolution in Western North America: A Case Study in Rosaceae. – PLoS one 7(12): e50358.

Veloz, S. D., Williams, J. W., Blois, J. L., He, F., Otto-Bliesner, B. and Liu, Z. 2012. No-analog climates and shifting realized niches during the late quaternary: implications for 21st-century predictions by species distribution models. – Global Change Biology 18(5): 1698–1713.

Venditti, C., Meade, A. and Pagel, M. 2011. Multiple routes to mammalian diversity. – Nature 479(7373): 393–396.

Verbruggen, H., Tyberghein, L., Pauly, K., Vlaeminck, C., Nieuwenhuyze, K., Kooistra, W., Leliaert, F. and Clerck, O. 2009. Macroecology meets macroevolution: evolutionary niche dynamics in the seaweed *Halimeda*. – Global Ecology and Biogeography 18(4): 393–405.

Vieites, D., Nieto-Román, S. and Wake, D. 2009. Reconstruction of the climate envelopes of salamanders and their evolution through time. – Proceedings of the National Academy of Sciences 106(Supplement 2): 19715–19722.

de Villemereuil, P., Wells, J. A., Edwards, R. D. and Blomberg, S. P. 2012. Bayesian models for comparative analysis integrating phylogenetic uncertainty. – BMC Evolutionary Biology 12(1): 102.

Webster, A. J. and Purvis, A. 2002. Ancestral states and evolutionary rates of continuous characters. – In: MacLeod, N. and Forey, P. L. (eds.), Morphology, shape and phylogeny. Taylor and Francis, London and New York, pp. 247–268.

Wiens, J., Kuczynski, C., Duellman, W. and Reeder, T. 2007. Loss and re-evolution of complex life cycles in marsupial frogs: does ancestral trait reconstruction mislead?. – Evolution 61(8): 1886–1899.

Wiens, J. J. 2008. Commentary on Losos (2008): niche conservatism deja vu. – Ecology Letters 11(10): 1004–1005.

Wiens, J. J., Ackerly, D. D., Allen, A. P., Anacker, B. L., Buckley, L. B., Cornell, H. V., Damschen, E. I., Jonathan Davies, T., Grytnes, J.-A., Harrison, S. P. et al. 2010. Niche conservatism as an emerging principle in ecology and conservation biology. – Ecology Letters 13(10): 1310–1324.

Wiens, J. J., Kozak, K. H. and Silva, N. 2013. Diversity and niche evolution along aridity gradients in North American lizards (Phrynosomatidae). – Evolution 67(6): 1715–1728.

Williams, J. W. and Jackson, S. T. 2007. Novel climates, no-analog communities, and ecological surprises. – Frontiers in Ecology and the Environment 5(9): 475–482.

Williams, J. W., Jackson, S. T. and Kutzbach, J. E. 2007. Projected distributions of novel and disappearing climates by 2100 AD. – Proceedings of the National Academy of Sciences 104(14): 5738–5742.

Williams, J. W., Shuman, B. N. and Webb III, T. 2001. Dissimilarity analyses of late-Quaternary vegetation and climate in eastern North America. – Ecology 82(12): 3346–3362.

Wright, S. 1934. An analysis of variability in number of digits in an inbred strain of guinea pigs. – Genetics 19(6): 506.

Yesson, C. and Culham, A. 2006a. Phyloclimatic modeling: combining phylogenetics and bioclimatic modeling. – Systematic Biology 55(5): 785–802.

Yesson, C. and Culham, A. 2006b. A phyloclimatic study of *Cyclamen*. – BMC Evolutionary Biology 6(1): 72.

